# Label-free visualization and quantification of the drug-type-dependent response of tumor spheroids by dynamic optical coherence tomography

**DOI:** 10.1101/2023.09.28.559912

**Authors:** Ibrahim Abd El-Sadek, Rion Morishita, Tomoko Mori, Shuichi Makita, Pradipta Mukherjee, Satoshi Matsusaka, Yoshiaki Yasuno

## Abstract

We demonstrate label-free dynamic optical coherence tomography (D-OCT)-based visualization and quantitative assessment of patterns of tumor spheroid response to three anti-cancer drugs. The study involved treating human breast adenocarcinoma (MCF-7 cell-line) with paclitaxel (PTX), tamoxifen citrate (TAM), and doxorubicin (DOX) at concentrations of 0 (control), 0.1, 1, and 10 μM for 1, 3, and 6 days. In addition, fluorescence microscopy imaging was performed for reference. The D-OCT imaging was performed using a custom-built OCT device. Two algorithms, namely logarithmic intensity variance (LIV) and late OCT correlation decay speed (OCDS_*l*_) were used to visualize the tissue dynamics. The spheroids treated with 0.1 and 1 μM TAM appeared similar to the control spheroid, whereas those treated with 10 μM TAM had significant structural corruption and decreasing LIV and OCDS_*l*_ over treatment time. The spheroids treated with PTX had decreasing volumes and decrease of LIV and OCDS_*l*_ signals over time at most PTX concentrations. The spheroids treated with DOX had decreasing and increasing volumes over time at DOX concentrations of 1 and 10 μM, respectively. Meanwhile, the LIV and OCDS_*l*_ signals decreased over treatment time at all DOX concentrations. The D-OCT, particularly OCDS_*l*_, patterns were consistent with the fluorescence microscopic patterns. The diversity in the structural and D-OCT results among the drug types and among the concentrations are explained by the mechanisms of the drugs. The presented results suggest that D-OCT is useful for evaluating the difference in the tumor spheroid response to different drugs and it can be a useful tool for anti-cancer drug testing.

## Introduction

Cancer is one of the most fatal diseases worldwide^1,2^, accounted for 10 million deaths in 2020^3^. The cancer mortality rate can be reduced through early diagnosis and the proper selection of anti-cancer drugs. There are several drug types that can be used for the treatment of a single type of cancer. As an example, approximately 80 drugs, including abemaciclib, paclitaxel (PTX), tamoxifen citrate (TAM), doxorubicin hydrochloride (DOX), and anastrozole, are approved for breast cancer treatment in the United States.

Each type of anti-cancer drug has a different mechanism of interaction with cancer cells. Abemaciclib, for example, is a cyclin-dependent kinase (CDK) inhibitor that reduces the proliferation of estrogen-receptor-positive breast cancers^4,5^. PTX corrupts the cell structure and halts cell mitosis by stabilizing microtubules(MTs)^6,7^. TAM is an estrogen receptor transcription inhibitor that up-regulates the production of transforming growth factor *β* (TGF-*β*) and down-regulates insulin-like growth factor 1 (IGF-1) and protein kinase C (PKC), resulting in cell apoptosis^8,9^. DOX inhibits the growth of breast cancer cells by blocking topoisomerase II and induces apoptosis^10,11^. Anastrozole, is an estrogen blocker^12^. Understanding the mechanisms of interaction between drugs and cancer cells is important for proper anti-cancer drug selection.

The tumor spheroid, which is a three-dimensional (3D) culture of cancer cells, can be used to clarify a drug-cell interaction^13,14^. A tumor spheroid closely emulates the heterogeneous structure, growth kinetics, and cell interactions of *in vivo* solid tumors^15^. The efficacy of an anti-cancer drug can be evaluated via the effect of the drug on the spheroid’s morphology and cell viability. The spheroid’s morphology and cell viability are assessed using several gold-standard modalities, such as histology^16,17^, fluorescence microscopy^18–20^, and bright-field microscopy^21,22^. However, these methods have several limitations. Staining histology is invasive and provides only two-dimensional slice imaging. Bright-field microscopy lacks molecular specificity and is a two-dimensional imaging method. Fluorescence microscopy is invasive and its deep imaging ability is limited to a few hundred microns. Meanwhile, the requirements for spheroid imaging are label-free, volumetric, deep, and longitudinal imaging capabilities.

In contrast with standard microscopy, optical coherence tomography (OCT) is a label-free and volumetric imaging modality having a long penetration depth, such as a depth of a few millimeters^23^. OCT is thus a useful microscopic imaging modality for tumor spheroid imaging. The necrotic regions of tumor spheroids were recently highlighted through OCT-based attenuation coefficient imaging^24,25^. In addition, the quantification of the volume and growth of a tumor spheroid was recently demonstrated using standard OCT imaging^26^. Furthermore, OCT-based tissue density quantification was used for evaluating regional differences in the drug response of tumor spheroids^27^. However, these methods quantify only the static properties of the tissue. To access the intracellular activity, which may reflect the cell viability, further extension of OCT is required.

Dynamic OCT (D-OCT) is an emerging functional extension of OCT that combines high-speed OCT systems and the signal processing of the rapidly captured OCT frames. D-OCT provides cellular and sub-cellular activity contrasts without using external contrast agents. Several studies have demonstrated the ability of D-OCT to visualize the intracellular activities of fresh *ex vivo* and *in vitro* samples including tumor spheroids^28–33^. Most of these methods are two-dimensional, i.e., they provide only *en face*^28^ or cross-sectional^30–32^images. Although volumetric D-OCT has been demonstrated, it requires long acquisition time or expensive very-high-speed OCT systems^29,33^. We recently proposed a 3D D-OCT method for imaging the intracellular dynamics of tumor spheroids by combining a standard-speed swept-source OCT device and a custom-designed 3D scanning protocol^34^. This method captured a 3D D-OCT volume in 52.4 s^34^. In our previous work, the viable and necrotic regions of tumor spheroids and their longitudinal alteration over 20 hours were successfully visualized by the 3D D-OCT method^34^. In addition, the 3D D-OCT was shown to be useful for visualizing and qualitatively analyzing specific drug-response patterns of human breast cancer (MCF-7 cell-line) and colon cancer (HT-29 cell-line) spheroids treated with PTX and the active metabolite of irinotecan (SN-38), respectively^35^.

In general, selecting the proper anti-cancer drug may require pre-clinical trials of multiple drugs for a patient-derived tumor spheroid. As different drugs have different mechanisms of interaction with tumor cells, the application of different drugs leads to different morphological and cell-activity-response patterns of the spheroids. To establish 3D D-OCT as a standard technique for drug effect evaluation, it is necessary to demonstrate the ability of 3D D-OCT to visualize and quantify the different responses of a tumor spheroid to various drug types.

In this paper, we demonstrate label-free assessment of tumor spheroid response to three types of anti-cancer drugs by 3D D-OCT. The study involves MCF-7 spheroids treated with TAM, PTX, and DOX. The drugs were applied at three concentrations for three administration times. We also measured non-treated spheroids as a control group. The D-OCT successfully showed that the response patterns were highly divergent for the three types of drugs. In addition, the morphological features and viability of the spheroids were quantified using the OCT images and D-OCT signals, and the time-course trends of these quantified metrics are shown. The time-course trends were also found to be drug/time specific. In the Discussion section, we show that these drug/time specific findings of OCT and D-OCT can be explained by the interaction mechanisms of the drugs.

## Results

OCT imaging was performed using a custom-made swept-source OCT microscope^36^. For the tumor spheroid dynamics imaging, we repeatedly captured 32 OCT frames at each location in the spheroid over a period of 6.348 s with an inter-frame interval of 204.8 ms. For the intracellular dynamics visualization, the temporal fluctuations of the sequentially captured dB-scaled OCT signal were analyzed using two D-OCT algorithms, namely logarithmic intensity variance (LIV) and OCT correlation decay speed (OCDS_*l*_)^30,34^. Further details of the OCT microscope and D-OCT algorithms are described in the Methods section.

### D-OCT imaging of control MCF-7 spheroids

The first row of images in Fig. 1 are pseudo time-course OCT intensity, LIV, OCDS_*l*_, and fluorescence images of the MCF-7 spheroids without drug administration (i.e., the controls). The OCT intensity images show a slight increase in the spheroid size over treatment time.

**Figure 1.**
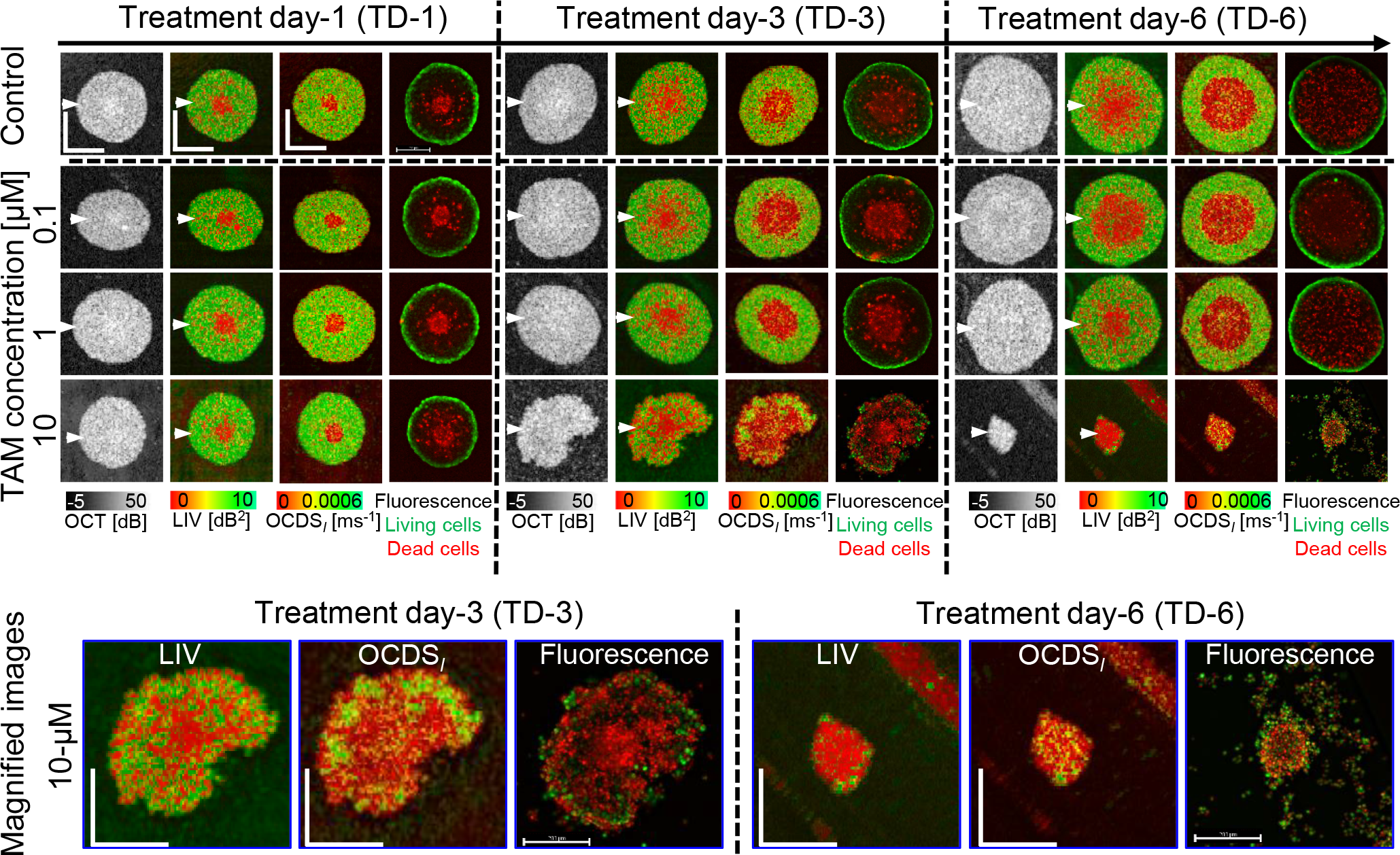
OCT intensity, LIV, OCDS_*l*_, and fluorescence images of the MCF-7 control and TAM-treated spheroids. The images are *en face* slices at approximately the center of the spheroids. The control spheroids (first row) show widening of the low LIV (red) and low OCDS_*l*_ (red) area at the center over time. The low LIV and OCDS_*l*_ are consistent with the dead cells concentrated at the spheroid center found in fluorescence images. The spheroids treated with 0.1 μM and 1μM TAM appear similar appearances to the control spheroids over time. In contrast, the spheroids treated with 10 μM show significant shape corruption, volume reduction, and decrease of D-OCT signals over treatment time. The scale bars represent 200 μm. The cross-sectional images of the same spheroids are available in the supplementary material (Fig. S1).

The D-OCT (i.e., LIV and OCDS_*l*_) images showed low LIV (red) and low OCDS_*l*_ (red) at the spheroid center. The low-signal region enlarged with increasing cultivation time. This D-OCT pattern is consistent with our previous studies^34,35^. The low LIV and OCDS_*l*_ at the spheroid center may correspond to dead cells at the spheroid center observed in the corresponding fluorescence images. MCF-7 is known to form a necrotic core at the spheroid center owing to a lack of oxygen and nutrients, whereas the spheroid periphery comprises viable tumor cells^37–39^. The low D-OCT at the spheroid center might indicate the necrotic core, whereas the high D-OCT at the spheroid periphery may indicate the viable tumor cells^30,34^.

### D-OCT imaging of TAM-treated MCF-7 spheroids

The second and later rows of Fig. 1 are the pseudo time-course images of the TAM-treated MCF-7 spheroids. At TAM concentrations of 0.1 and 1 μM, the OCT intensity images show a slight increase in the spheroid size over the treatment time (Fig. 1, second and third rows). At a TAM concentration of 10 μM (fourth row), structural corruptions are evident after long treatment times (treatment day (TD)-3 and -6).

D-OCT images (LIV and OCDS_*l*_) for TAM concentrations of 0.1 and 1 μM (second and third rows) have patterns similar to those of the control (i.e., untreated) spheroids. In addition, the fluorescence images have patterns similar to those of the control spheroids. This suggests that TAM is not effectual at concentrations of 0.1 and 1 μM. In contrast, spheroids treated with 10 μM (fourth row) of TAM have high occupancy of low-LIV and low-OCDS_*l*_ regions, which become larger for the longer treatment time. Magnified images of representative cases of severe structural corruption, size reduction, and reduction of LIV and OCDS_*l*_ signals are shown at the bottom of the figure. The low LIV and low OCDS_*l*_ signals are found to coincide with the appearance of dead cells (red) in the fluorescence images. This coincidence and the structural corruption indicate the high efficacy of TAM at a concentration of 10 μM. The diverse response of the spheroid among the concentrations of TAM can be explained by the drug-cell interaction mechanism of TAM as discussed in detail in the Discussion section.

The cross-sectional OCT intensity and D-OCT images extracted at the locations indicated by white arrowheads in Fig. 1 have tendencies similar to those of the *en face* images in Fig. 1 for both control and TAM cases. (See Supplementary Fig. S1). In addition, the results of two additional spheroids measured at each treatment condition of TAM and the control show similar findings to those presented in Fig. 1 (supplementary Figs. S2 and S3).

The spheroid volume, mean LIV, mean OCDS_*l*_, and LIV- and OCDS_*l*_-based necrotic cell ratios (see the Method section) of the control and TAM-treated spheroids at each time point are shown in Fig. 2. The volumes of the spheroids treated with 0.1 and 1 μM TAM increase monotonically with time, whereas the volume of the spheroid treated with 10 μM TAM significantly decreases on TD-6 [Fig. 2 (a)]. The mean LIV and mean OCDS_*l*_ [Fig. 2 (b) and (c), respectively] have decreasing trends at all concentrations including the control. Among all the concentrations, 10 μM of TAM shows the most pronounced reduction in mean LIV and mean OCDS_*l*_. At TAM concentrations of 0.1 and 1 μM, the LIV- and OCDS_*l*_-based dead cell ratios [Fig. 2 (d) and (e), respectively] increase slightly over time. In contrast, more rapid and larger increases in the dead cell ratio are observed at 10 μM of TAM. It is noted that all plots for TAM concentrations of 0.1 μM and 1 μM are remarkably similar to those of the control, whereas the plots of 10 μM exhibits more pronounced and distinctive trends.

**Figure 2.**
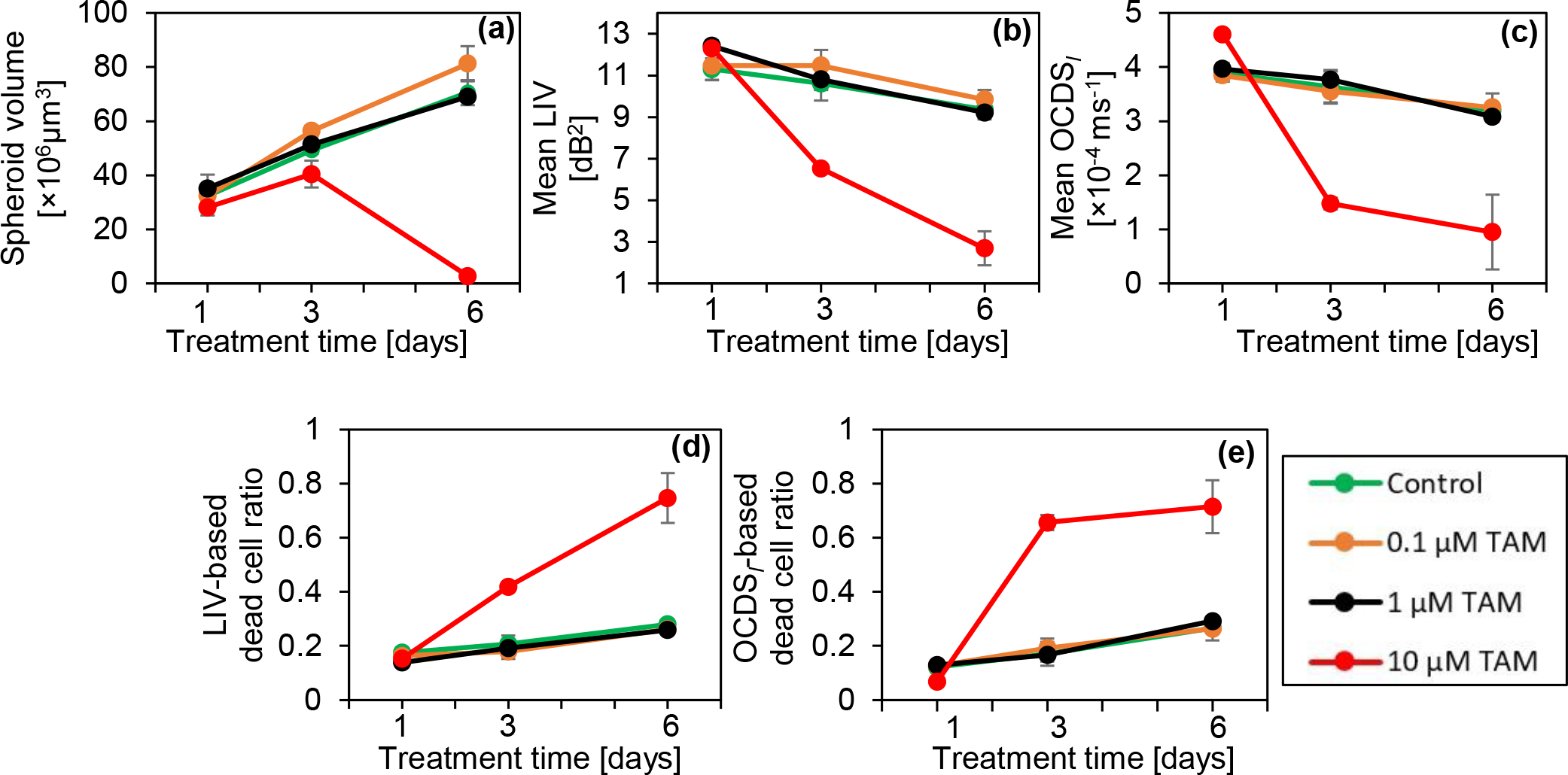
Morphological and D-OCT signal quantification of the control and TAM-treated MCF-7 spheroids. The plotted values are averages for three spheroids measured under each condition, whereas the error bars represent the ± standard-deviation range. The 0.1 and 1 μM cases show similar results to those in the control case, whereas the 10 μM case shows significant departure.

### D-OCT imaging of PTX-treated MCF-7 spheroids

Figure 3 presents the OCT, D-OCT, and fluorescence images of MCF-7 spheroids treated with PTX. The OCT images of 1 and 10 μM PTX show a slight reduction in the spheroid size over time, whereas the spheroid size increases slightly for treatment with 0.1 μM PTX. Besides the central low-LIV (red) core, low-LIV (red) spots are observed for all concentrations, and the size and number of the low-LIV spots increase over time (first to third rows). Meanwhile, there are several low-OCDS_*l*_ domains rather than spots (indicated by arrow heads), which are not observed in the LIV. Magnified images (bottom row) show that these low-OCDS_*l*_ domains are more continuous and larger at a higher concentration of PTX. In addition, for treatment with 1 and 10 μM PTX, the necrotic cores, the red central regions in the LIV and OCDS_*l*_, found on TD-1 and -3 disappear on TD-6. The fluorescence images show an increase in the dead cell area (red) over treatment time at all PTX concentrations, which appears consistent with the increase of low LIV and low OCDS_*l*_ regions.

**Figure 3.**
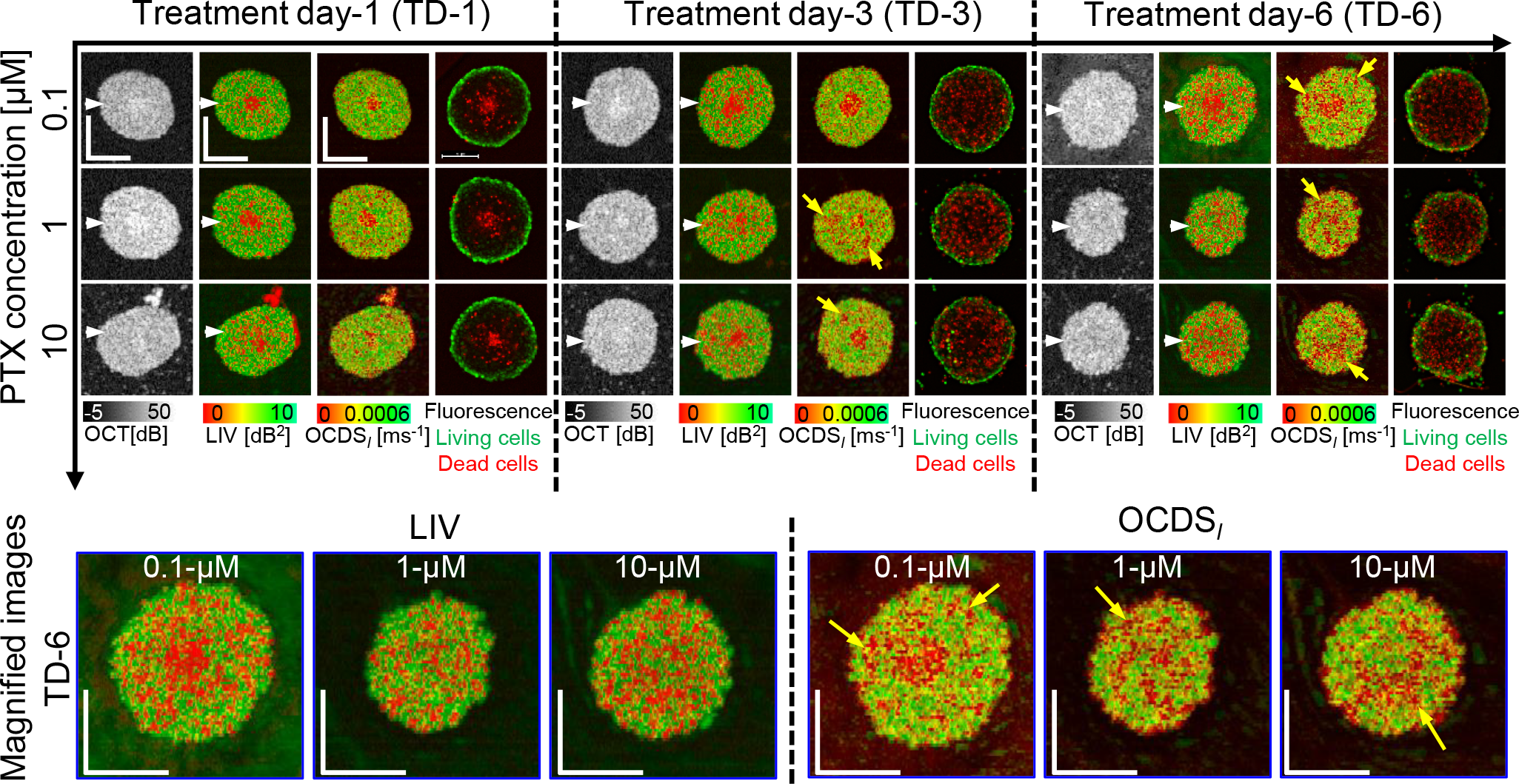
The *en face* OCT intensity, LIV, OCDS_*l*_, and fluorescence images of the MCF-7 spheroids treated with PTX. OCT intensity images show moderate reduction of the spheroid size over time at all PTX concentrations. The low-LIV and low-OCDS_*l*_ regions (red regions) increase over treatment time at all PTX concentrations. Fluorescence images show an increase in the number of dead cells (red, stained by PI) over time at all PTX concentrations. The scale bars represent 200 μm. The cross-sectional images of the same spheroids are available in the supplementary material (Fig. S4).

The cross-sectional OCT and D-OCT images extracted at the location of the white arrow heads in Fig. 3 are summarized in the supplementary material (Fig. S4) and have similar appearances. In addition, the results for two additional MCF-7 spheroids measured under each PTX treatment condition are summarized in supplementary material (Figs. S5 and S6). The spheroids appear similar to those presented in Fig. 3.

Figure 4 shows the quantitative analysis of PTX-treated spheroids similar to Fig. 2. The plots of the control spheroids are reprinted in the figure. At all concentrations, the spheroid volume becomes markedly smaller than the control [Fig. 4(a)]. In particular, the spheroids treated with 1 and 10 μM PTX have moderate volume reductions, whereas the spheroid treated with 0.1 μM PTX has a slight increase. The mean LIV and mean OCDS_*l*_ [Fig. 4 (b) and (c), respectively] monotonically decrease over time at all PTX concentrations. This decreasing tendency is similar to that of control, but relatively more pronounced than the control. The LIV- and OCDS_*l*_-based dead cell ratios [Fig. 4 (d) and (e)] increase monotonically over time at all PTX concentrations, which is similar to that of control but relatively more pronounced.

**Figure 4.**
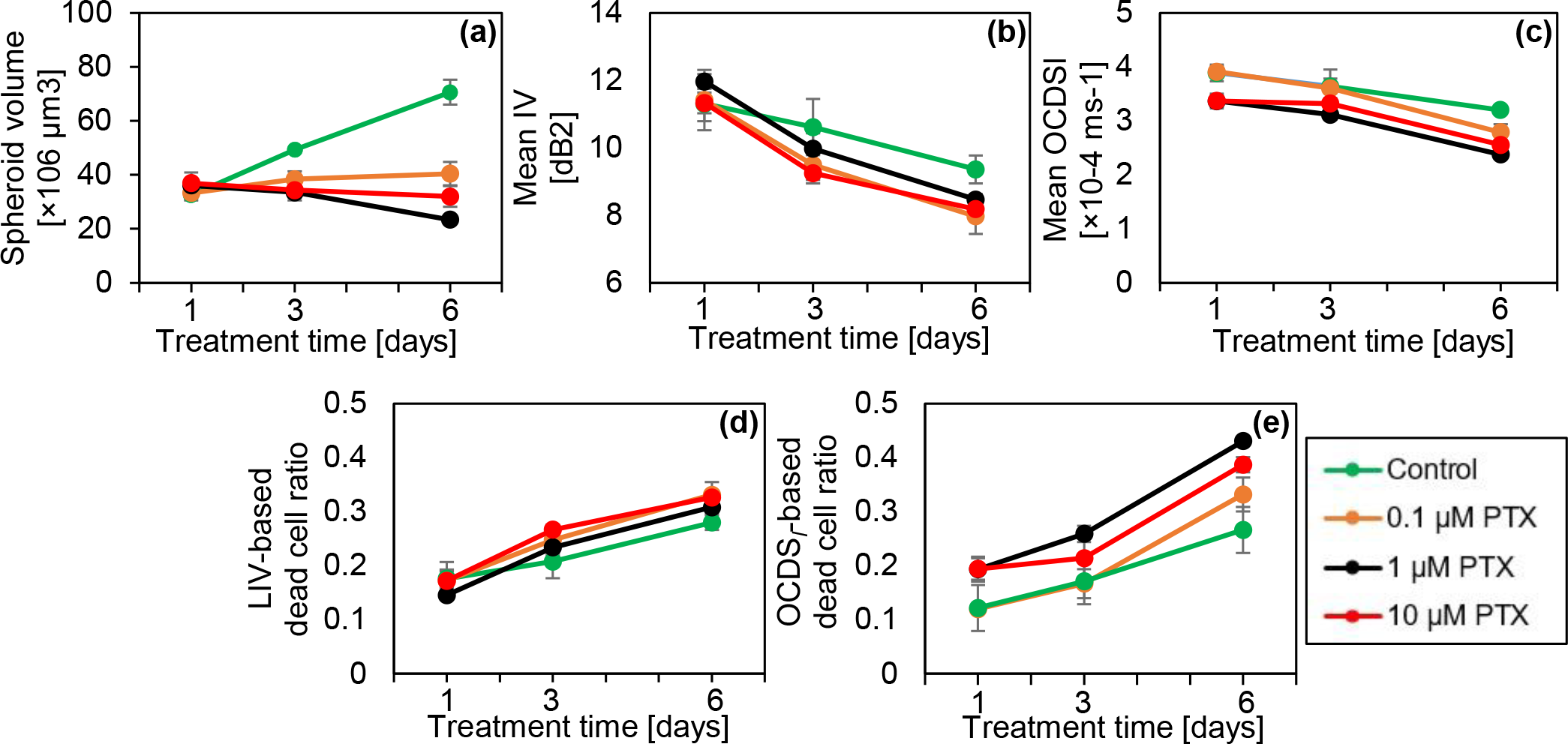
Quantification of the OCT and D-OCT of MCF-7 spheroids treated with PTX. The figures are displayed in the same order with Fig. 2.

The reduction in the spheroid volume found in the images [Fig. 3] and plot [Fig. 4(a)] may suggest the corruption of the cells’ structure and shape under the PTX application. Meanwhile, the decreases in the mean LIV and mean OCDS_*l*_ [Fig. 4(b) and (c)] and the increase in the number of low-LIV spots and low-OCDS_*l*_ domains may suggest the PTX-induced reduction of intracellular transport through MTs. These points are discussed in more detail in the Discussion section.

### D-OCT imaging of the DOX-treated MCF-7 spheroids

Figure 5 presents the results for the MCF-7 spheroids treated with DOX. For 0.1 μM [Fig. 5 (first row)], following features were found. Namely, the OCT images show that the spheroid surface becomes hazy after a long treatment time. The LIV images show that the number of low-LIV spots increases over time and the necrotic core, which is the centermost red region, disappears on TD-6. The OCDS_*l*_ images have similar patterns on TD-1 and TD-3, whereas there are several low-OCDS_*l*_ domains on TD-6, which are not clear in the corresponding LIV images. The fluorescence images in the 0.1-μM case show an increase in the number of dead cells over treatment time, which appear consistent with the increase in the number of low-LIV spots and low-OCDS_*l*_ domains over time.

**Figure 5.**
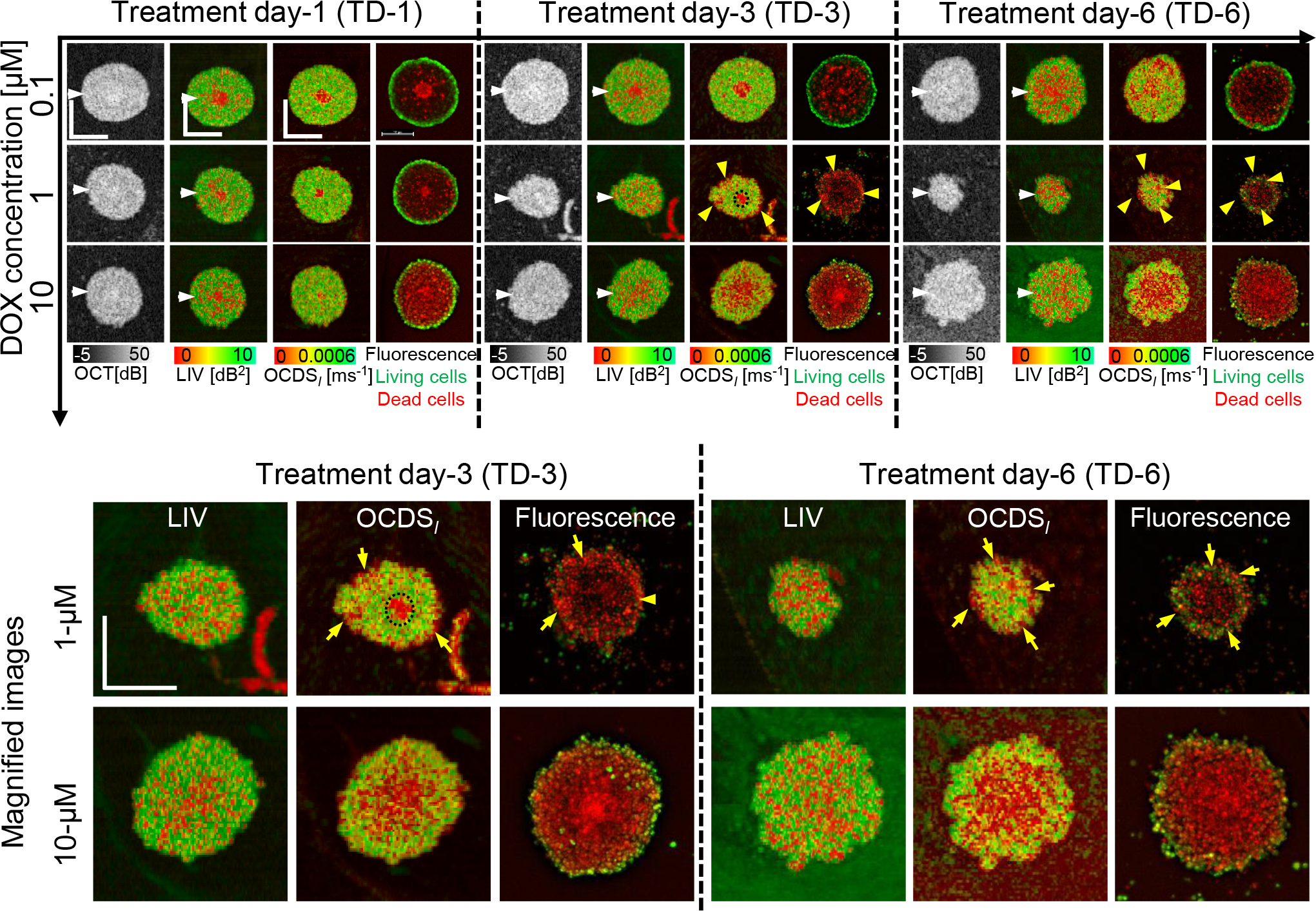
The *en face* OCT intensity, LIV, OCDS_*l*_, and fluorescence images of the MCF-7 spheroids treated with DOX. Shrinkage at a treatment concentration of 1 μM and swelling at 10 μM are observed in OCT. At 0.1 μM, the number of low-LIV (red) spots increases over time, whereas OCDS_l_ shows domains of low OCDS_l_ (red) on TD-6. The spheroid treated with 1 μM DOX shrinks over time and has thin low-OCDS_l_ regions near the outer surface, which are consistent with dead cells in fluorescence images. In the 10-μM case, the spheroids swell over time. The OCDS_l_ patterns are consistent with the fluorescence images. The scale bars represent 200 μm. Cross-sectional images of the same spheroids are presented in Fig. S7 in the supplementary material. The scale bars represent 200 μm.

In the case of 1 μM DOX [Fig. 5 (second row)], the spheroid markedly shrinks over time. The low-LIV spots increase over time and become dominant on TD-6. In contrast, the OCDS_*l*_ images show thin low-OCDS_*l*_ regions at the spheroid periphery (yellow arrowheads) and a circular low-OCDS_*l*_ core at the center on TD-3, which are not observed in LIV as shown in the magnified images (first row of the magnified images). The low OCDS_*l*_ at the spheroid periphery may indicate the interaction of DOX with the most outer layer of the spheroid, whereas the circular low OCDS_*l*_ at the center may indicate the necrotic core. The low-OCDS_*l*_ core disappears on TD-6. The peripheral low-OCDS_*l*_ regions observed on TD-3 and TD-6 (yellow arrowheads) are consistent with the dead cells, concentrated around the spheroid periphery in the fluorescence images (yellow arrowheads). It may indicate the tumor cell death at the spheroid periphery induced by DOX.

In contrast with the 1 μM case, the spheroids treated with 10 μM DOX (Fig. 5, third row) swell over time. The spheroid shrinkage and swelling at 1 and 10 μM DOX, respectively may indicate different mechanisms of DOX as discussed in the Discussion section. The LIV images for 10 μM DOX show granular patterns of low and high LIV, and the low-LIV spots become ubiquitous on TD-6. Meanwhile, the OCDS_*l*_ shows concentric appearances, where a low-OCDS_*l*_ region is surrounded by high-OCDS_*l*_ as shown in the second row of magnified images (Fig. 5, fourth and fifth rows). The low and high OCDS_*l*_ layers are consistent with dead and living cell patterns in the fluorescence images more than LIV.

The cross-sectional OCT and D-OCT images of the MCF-7 spheroids treated with DOX are summarized in the supplementary material (Fig. S7). In addition, two additional spheroids measured under each treatment condition with DOX have response patterns similar to those presented in Fig. 5. (See supplementary Figs. S8 and S9.)

Figure 6 presents the quantitative analysis of the DOX-treated spheroids similar to Fig. 2. In comparison with the control, all spheroids treated with DOX show a markedly smaller volume [Fig. 6(a)]. In the treatment with 0.1 μM DOX, the spheroid volume decreases over time, whereas in the treatment with 1 μM DOX, the spheroid has marked decreasing trend that corresponds to the shrinkage observed in Fig. 5. In contrast, the volume of the spheroid treated with 10 μM DOX increases slightly over time, which may correspond to the spheroid swelling observed in Fig. 5. The mean LIV [Fig. 6(b)] and mean OCDS_*l*_ [Fig. 6(c)] decrease over time at all DOX concentrations. Meanwhile, the LIV- and OCDS_*l*_-based dead cell ratios [Fig. 6(d) and (e), respectively] increase over time at all DOX concentrations. We suspect that the reduction of mean D-OCT signals and increase of dead cell ratios indicate cell apoptosis induced by DOX as discussed in detail in the Discussion section.

**Figure 6.**
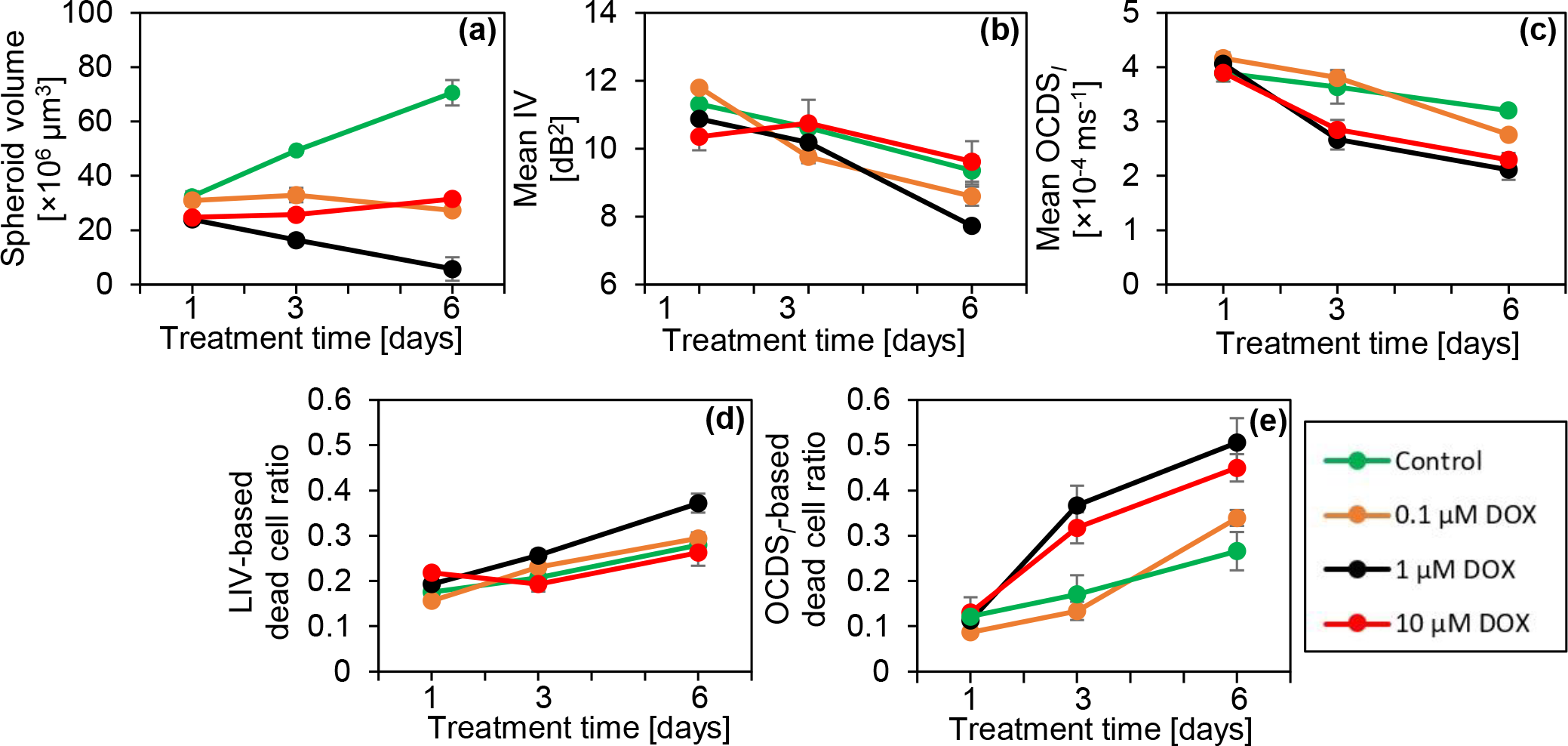
Quantification of the MCF-7 spheroids treated with DOX. The figures are displayed in the same order with Figs. 2 and 4. For the 1 μM case, the spheroid volume significantly decreases over time, whereas it increases over time for the 10 μM case. This drug-concentration dependency can be attributed to the interaction mechanisms of DOX. (See the Discussion for details.)

## Discussion

For the 0.1 and 1 μM TAM cases, the D-OCT images showed similar image patterns to those in the control cases (Fig. 1). In addition, these cases showed similar increasing trends of the spheroid volume to those in the control cases [Fig. 2 (a)]; i.e., TAM is ineffectual at these concentrations. In contrast, for the 10-μM case, significant shape corruption, volume reduction, and reduction of the D-OCT signals were observed. The difference in the MCF-7 spheroid responses to the low (0.1 and 1 μM) and high (10 μM) concentrations of TAM can be explained by the mechanism of action of TAM as follows.

TAM is known to have two mechanisms of drug-cell interactions^8^. In the first mechanism, the TAM behaves as an estrogen-receptor (ER) antagonist; i.e., TAM inhibits ER transcription^8,40^. The ER is a female hormone located in the cell and is activated by estrogen. Following activation, ERs are translocated within the cell’s nucleus and work as a DNA-binding transcription factor^41^. The inhibition of ER thus inhibits the proliferation of ER-positive breast cancer cells. In the second mechanism, TAM directly or indirectly inhibits PKC. The inhibition is direct for a high concentration of TAM and indirect for a low concentration. The PKC inhibition causes apoptosis and finally inhibits cellular proliferation^8,9^. This mechanism can be effectual for ER-negative breast cancer.

As our MCF-7 spheroid is not co-cultured with estrogen-supporting cells, the first mechanism cannot be effectual. This can explain why the OCT and D-OCT findings of 0.1 and 1 μM TAM cases are nearly identical to those in the control case. In addition, we suspect that the indirect PKC inhibitions are not effectual in these cases because of the cells’ resistance to TAM^42^. In contrast, at a concentration of 10 μM, TAM may act as the direct inhibitor of PKC; i.e., the second mechanism of high concentration. The shape corruption and volume reduction observed by OCT may indicate the stopping of cell proliferation, whereas the low D-OCT signals observed at 10 μM TAM may indicate the cell apoptosis induced by the second mechanism of TAM. Furthermore, it has been reported that 10 μM TAM completely killed tumor cells in MCF-7 spheroid in one week^43^, which is consistent with our findings.

The structural and dynamic OCT findings of the MCF-7 spheroids treated with PTX [Fig. 3] can be explained by the interaction mechanism of PTX. PTX is known as an MT stabilizing drug used for breast cancer treatment. An MT is a polymer of tubulin dimers and a main constituent of the cytoskeleton^44^. The MT plays three vital roles in the cellular processes. First, as one of the main constituents of the cytoskeleton, MTs maintain the cell structure and support the cell with mechanical resistance against deformation^45^. Second, MTs play an important role in mitosis cell division^44,46^. Third, MTs form a platform for intracellular vesicles and organelles transport^44,47^. To fulfill these functions, MTs are highly dynamic. Namely, they grow and shrink rapidly and stochastically^45^, which allows them to bound to intracellular components, e.g., capturing chromosomes during mitosis.

The application of PTX prevents MT depolymerization and stabilizes the MT dynamic behavior^6,7^, resulting in cell-cycle and mitosis arrest followed by cell death. The reduction of LIV over time observed at all PTX concentrations may indicate a decrease in the magnitude of intracellular transport through MTs. Meanwhile, the reduction of OCDS_*l*_ may indicate the reduction of the speed of intracellular transport through MTs. Moreover, the reduction of the MCF-7 spheroid volume over time at 1 and 10 μM [Fig. 4(a)] may indicate the inhibition of cell mitosis due to the stabilization of MTs induced by PTX.

The DOX-treated spheroids shrunk at a treatment concentration of 1 μM and swelled at a concentration of 10 μM [Figs. 5 and 6(a)], and D-OCT signals decreased over the treatment time at all concentrations [Fig. 6(b) and (c)]. These findings can be explained by the mechanism of DOX as follows. DOX is known to inhibit the growth of cancer cells by blocking the enzyme topoisomerase II, which is responsible for DNA repair^10,11^. This inhibition results in cell apoptosis. In addition, DOX is responsible for releasing free radicals that damage cell membranes^10^. The shrinkage of the spheroid at 1 μM of DOX may indicate the growth inhibition of the tumor cells. Meanwhile, spheroid swelling at 10 μM may indicate the cell membrane damage and the loosening of the bonds between cells. Moreover, the reduction of the LIV and OCDS_*l*_ signals over time may indicate the apoptosis induced by DOX application.

In the present study, the LIV and OCDS_*l*_ images showed different patterns. For example, the OCDS_*l*_ images of the control, and 0.1 and 1 μM TAM spheroids showed clearer boundaries of the necrotic core than the LIV images [Fig. 1 (first to third rows)]. In addition, in the treatments of 10 μM TAM on TD-3 [Fig. 1 (fourth row)], all PTX concentrations on TD-6 [Fig. 3], and 10 μM DOX on TD-3 and TD-6 [Fig. 5 (4th row)], the LIV images show granular patterns of low and high signals, while the OCDS_*l*_ show domain structures. In principle, LIV is supposed to be a measure of the magnitude of intracellular motility, whereas OCDS_*l*_ is a measure of the speed of intracellular motility^30,34^. It is thus expected to obtain different image patterns for LIV and OCDS_*l*_ and we believe that the two methods complement each other. A variety of intracellular activities are known to occur on different time scales^48^. Hence, speed metrics of the intracellular motility, such as OCDS_*l*_, can be a more specific marker to a particular intracellular activity than LIV. Meanwhile, LIV is a metric that can be used directly to visualize the magnitude of intracellular activity.

## Conclusion

We demonstrated the ability of D-OCT to visualize and quantitatively assess different patterns of response of tumor spheroids to different anti-cancer drugs. Three anti-cancer drugs were applied to human breast adenocarcinoma (MCF-7 cell-line) spheroids, and D-OCT showed different morphological and tissue dynamics patterns. The drug-type-dependent alterations of the spheroid morphology and intracellular dynamics were successfully visualized and quantified by the 3D D-OCT method. The drug-type-dependent variations in the D-OCT patterns were suggested to indicate the different mechanisms of anti-cancer drugs. In conclusion, D-OCT can be used to visualize and quantify the different responses of tumor spheroids to different anti-cancer drugs and it can be a useful tool for the assessment of anti-cancer drugs.

## Methods

### Spheroid preparation and drug treatment protocols

Figure 7 is a time chart schematic of the tumor spheroid drug response study design. On day-1, 1000 cells derived from human breast cancer (MCF-7 cell-line) and purchased from the Japanese Collection of Research Bioresources cell bank were seeded in each well of an ultra-low-attachment 96-well-plate. We prepared 90 wells in total. The cells were kept in the cell culture environment at a temperature of 37 °C and with a 5% CO_2_ supply. The cell-culture medium contained a 1:1 mixture of Eagle’s minimal essential medium (EMEM; for MCF-7) and F12 (Invitrogen, Waltham, MA) with a 2% B-27 supplement (Invitrogen), 2 ng/mL of basic fibroblast growth factor (bFGF; Wako, Osaka, Japan), 2 ng/mL of epidermal growth factor (EGF; Sigma-Aldrich, St. Louis, MO), 100 U/mL of penicillin G, and 0.1 mg/mL of streptomycin sulfate (Wako, Osaka, Japan). The cells aggregated with each other and formed an MCF-7 spheroid in each well on day-5.

**Figure 7.**
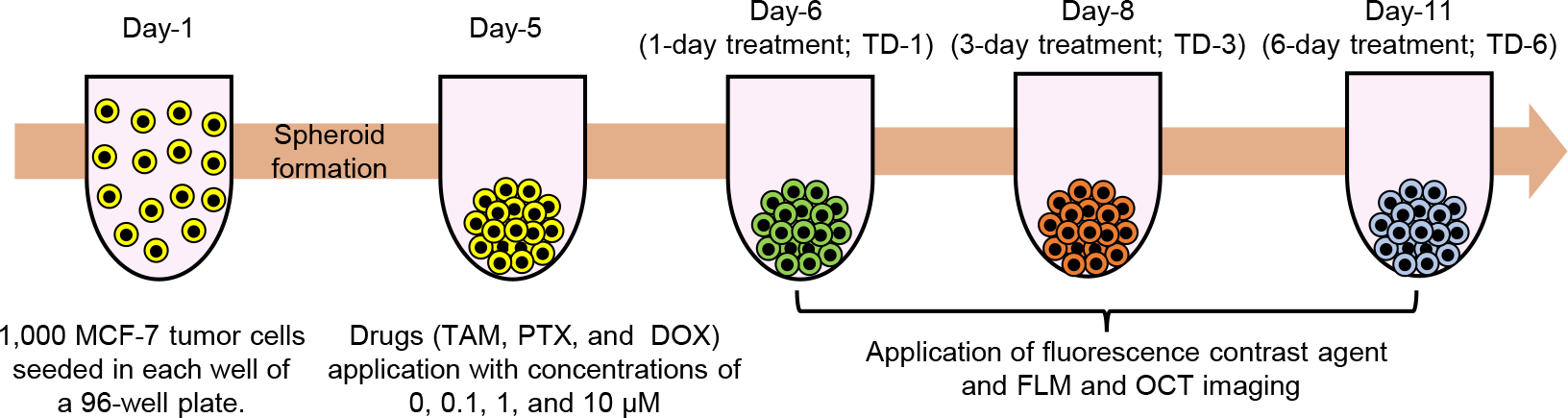
Time chart of the MCF-7 spheroid cultivation and measurement. In total, 90 spheroids were cultured. Each spheroid was formed by seeding 1,000 cells on day-1. The drugs, namely PTX, TAM, and DOX, were applied on day-5. On days-6, 8, and 11, fluorescence and then OCT imaging were performed. The fluorescence contrast agents (i.e., calcein-AM and PI) were applied for 3 hours before the fluorescence imaging.

Three anti-cancer drugs, namely TAM, PTX, and DOX, were applied on day-5 at concentrations of 0.1, 1, and 10 μM for each drug. Each spheroid was treated with one drug type at one concentration. In addition, untreated spheroids, denoted by a concentration of 0 μM, were kept as a control group for comparison. After drug application, the spheroids were kept in the same culture environment. The spheroids were then extracted from the cultivation and measured using the fluorescence and D-OCT microscope at three treatment time points denoted as treatment days-1, -3 and -6 (TD-1, TD-3, and TD-6). At each time-point, three spheroids were measured for each of the three drug concentrations and each of the three drug types. In addition, three control spheroids were measured at each treatment time point. Hence, 30 spheroids were measured at each of the three treatment time points, giving a total of 90 spheroids. It should be noted that, even for the same drug type and concentration, the spheroids measured at each time point were different individuals, as indicated by different spheroid colors on the time chart (Fig. 7). Hence, this study was not actually longitudinal but rather cross-sectional. We selected the cross-sectional design, because the D-OCT imaging was performed at a room temperature of approximately 25 °C without CO_2_ supply. The spheroids were thus no longer useful after the measurement owing to the lack of cultivation conditions. The integration of the D-OCT system with a cultivation environment may enable future longitudinal study.

### Fluorescence microscopic imaging

Fluorescence microscopy imaging was performed as a reference gold-standard method for tissue viability visualization. Two fluorescence agents were applied 3 hours before the fluorescence imaging. The first was calcein-acetoxymethyl (calcein-AM; Dojindo, Kumamoto, Japan), which labels living and/or viable cells and emits green fluorescence. The second agent was propidium iodide (PI; Dojindo), which labels dead cells and emits red fluorescence. The fluorescence imaging was performed using a wide-field fluorescence microscope (THUNDER imager DMi8; Leica Microsystems, Wetzlar, Germany) featuring a microscopic objective lens with a numerical aperture of 0.12.

### OCT device

A custom-built swept-source OCT device was used in this study. A microelectromechanical system (MEMS)-based swept source light source (AXP50124-8, Axun Technologies, MA) with a central wavelength of 1.31 μm and a scanning speed of 50,000 A-lines/s was used. A scan lens (LSM03, Thorlabs Inc., NJ) with an effective focal length of 36 mm was used. The lateral and axial (in tissue) resolutions were 18.1 μm and 14 μm, respectively. The power incident on the sample was 17 mW. More details about the OCT device have been published elsewhere^36^.

### Dynamic optical coherence tomography (D-OCT) imaging

#### 3D D-OCT scanning protocol

For 3D D-OCT imaging, we used our previously designed 3D scanning protocol^34^. Using this protocol, the lateral field of view of 1 ×1 mm^2^ was divided into eight blocks with each block containing 16 B-scan locations. Each block was scanned using a rapid raster scanning pattern and the raster scan was repeated 32 times in 6.348 s; i.e., at each B-scan location, 32 cross-sectional frames were captured as a time sequence with a frame repeating time of 204.8 ms. The volumetric data comprising 4096 frames captured at 128 B-scan locations in the spheroid were acquired in 52.4 s.

#### Logarithmic intensity variance (LIV) algorithm

The LIV is the time variance of the dB-scaled OCT. LIV measures the magnitude of dB-scaled OCT intensity fluctuations and is supposed to be sensitive to the magnitude of the intracellular motility^30,34^. By applying the LIV computation algorithm to the OCT data acquired by the above-mentioned protocol, a raw LIV volume comprising 128 B-scan locations was obtained. The final pseudo-color LIV image was a composition of the raw LIV and OCT intensity. The hue and value (brightness) of the image were proportional to the LIV and the average OCT intensity, respectively, and a maximum saturation of 1 was set for all pixels. Further mathematical descriptions of LIV are given in Refs.^30,34^.

#### Late OCT correlation decay speed (OCDS_l_) algorithm

To compute the OCDS_*l*_, we first computed the temporal auto-correlation of the OCT time sequence at each time delay point, so that we defined a decay curve of the auto-correlation as a function of the delay time. The OCDS_*l*_ was then computed as the slope of the auto-correlation decay curve at a certain delay range of [204.8, 1228.8 ms]. As OCDS_*l*_ quantifies the time characteristics of the sequentially captured OCT signal, it is supposed to be sensitive to the speed of the intracellular motility^30,34^. A pseudo color image of OCDS_*l*_ was generated in the similar manner to the LIV except for the hue channel, which was proportional to the OCDS_*l*_ value in this case. Refer to Refs. [^30,34^] for detailed mathematical descriptions.

#### Volumetric quantification of morphological and D-OCT signals

A two-step semi-automatic segmentation method was developed to quantify the morphological and dynamics signals of the tumor spheroids. In the first step, the spheroid segmentation was performed B-scan by B-scan using the find-connected-region plugin of Fiji ImageJ software with a manually defined OCT intensity threshold^49^. The segmentation mask generated in this first step contained the reflection from the bottom surface of the 96-well plate. In the second step, the well plate surface was manually removed B-scan by B-scan. Finally, the region containing only the spheroid tissue was segmented. Using this segmentation, the spheroid volume, and mean values of the LIV and OCDS_*l*_ of the entire spheroid volume were computed.

Furthermore, by applying empirically defined cut-offs of 3 dB^2^ and 2× 10^−4^ ms^−1^ for the LIV and OCDS_*l*_ respectively, the dead cell volume (i.e., the total volume of voxels with LIV or OCDS_*l*_ values lower than the cut-off) was computed. The dead cell ratio was then computed as the dead cells volume divided by the entire spheroid volume. The computed quantities were plotted as a function of the treatment time for each drug type and concentration, as shown in the Results section.

## Acknowledgments

Funding was provided by Core Research for Evolutional Science and Technology (JPMJCR2105), and the Japan Society for the Promotion of Science (JSPS) through the grants 18H01893, 21H01836, 22K04962, 22KF0058, and 22F22355. The first author is supported by the Postdoctoral Fellowships for Research in Japan program of JSPS (22F22355).

The authors greatly appreciate fruitful technical discussions held with Arata Miyazawa (Sky Technology).

## Author contributions statement

I.A., T.M., S.M., and Y.Y. designed the study. T.M. and S.M. prepared the samples. I.A. and R.M. organized the OCT experiments and collected all the data. I.A., R.M., Sh.M., P.M., S.M., and Y.Y. analyzed and interpreted the data. I.A. wrote the first draft of the paper, and all the authors revised the work and approved the final version of the manuscript. Y.Y. supervised the work.

## Competing interests

Abd El-Sadek, Makita, Mukherjee, Yasuno: Yokogawa Electric Corp. (F), Sky Technology (F), Nikon (F), Kao Corp. (F), Topcon (F). Mori, Matsusaka: None.

## Additional Information

Additional information can be found in the supplementary material.

## Data availability

Correspondence and requests for materials should be addressed to Y.Y.

## References

1. Sung, H. et al. Global Cancer Statistics 2020: GLOBOCAN Estimates of Incidence and Mortality Worldwide for 36 Cancers in 185 Countries. CA: A Cancer J. for Clin. 71, 209–249, DOI: 10.3322/caac.21660 (2021).

2. Bray, F., Laversanne, M., Weiderpass, E. & Soerjomataram, I. The ever-increasing importance of cancer as a leading cause of premature death worldwide. Cancer 127, 3029–3030, DOI: 10.1002/cncr.33587 (2021).

3. Chhikara, B. S. & Parang, K. Global Cancer Statistics 2022: the trends projection analysis. Chem. Biol. Lett. 10, 451, DOI: https://scholar.google.com/scholar?q=urn:nbn:sciencein.cbl.2023.v10.451 (2022).

4. Goetz, M. P. et al. MONARCH 3: A randomized phase III study of anastrozole or letrozole plus abemaciclib, a CDK4/6 inhibitor, or placebo in first-line treatment of women with HR+, HER2-locoregionally recurrent or metastatic breast cancer (MBC). J. Clin. Oncol. 33, TPS624–TPS624, DOI: 10.1200/jco.2015.33.15_suppl.tps624 (2015).

5. Kwapisz, D. Cyclin-dependent kinase 4/6 inhibitors in breast cancer: palbociclib, ribociclib, and abemaciclib. Breast Cancer Res. Treat. 166, 41–54, DOI: 10.1007/s10549-017-4385-3 (2017).

6. Xiao, H. et al. Insights into the mechanism of microtubule stabilization by Taxol. Proc. Natl. Acad. Sci. 103, 10166–10173, DOI: 10.1073/pnas.0603704103 (2006).

7. Weaver, B. A. How Taxol/paclitaxel kills cancer cells. Mol. Biol. Cell 25, 2677–2681, DOI: 10.1091/mbc.e14-04-0916 (2014).

8. Jordan, V. C. A current view of tamoxifen for the treatment and prevention of breast cancer. Br. J. Pharmacol. 110, 507–517, DOI: 10.1111/j.1476-5381.1993.tb13840.x (1993).

9. Radin, D. P. & Patel, P. Delineating the molecular mechanisms of tamoxifen’s oncolytic actions in estrogen receptor-negative cancers. Eur. J. Pharmacol. 781, 173–180, DOI: 10.1016/j.ejphar.2016.04.017 (2016).

10. Pilco-Ferreto, N. & Calaf, G. M. Influence of doxorubicin on apoptosis and oxidative stress in breast cancer cell lines. Int. J. Oncol. 49, 753–762, DOI: 10.3892/ijo.2016.3558 (2016).

11. Pengnam, S. et al. Synergistic Effect of Doxorubicin and siRNA-Mediated Silencing of Mcl-1 Using Cationic Niosomes against 3D MCF-7 Spheroids. Pharmaceutics 13, 550, DOI: 10.3390/pharmaceutics13040550 (2021).

12. Milani, M., Jha, G. & Potter, D. A. Anastrozole Use in Early Stage Breast Cancer of Post-Menopausal Women. Clin. medicine. Ther. 1, 141–156 (2009).

13. Yuhas, J. M., Li, A. P., Martinez, A. O. & Ladman, A. J. A Simplified Method for Production and Growth of Multicellular Tumor Spheroids. Cancer Res. 37, 3639–3643 (1977).

14. Hirschhaeuser, F. et al. Multicellular tumor spheroids: An underestimated tool is catching up again. J. Biotechnol. 148, 3–15, DOI: 10.1016/j.jbiotec.2010.01.012 (2010).

15. Costa, E. C. et al. 3D tumor spheroids: an overview on the tools and techniques used for their analysis. Biotechnol. Adv. 34, 1427–1441, DOI: 10.1016/j.biotechadv.2016.11.002 (2016).

16. Gaskell, H. et al. Characterization of a functional C3A liver spheroid model. Toxicol. Res. 5, 1053–1065, DOI: 10.1039/c6tx00101g (2016).

17. Jeppesen, M. et al. Short-term spheroid culture of primary colorectal cancer cells as an in vitro model for personalizing cancer medicine. PLOS ONE 12, e0183074, DOI: 10.1371/journal.pone.0183074 (2017).

18. Galateanu, B. et al. Impact of multicellular tumor spheroids as an in vivo-like tumor model on anticancer drug response. Int. J. Oncol. 48, 2295–2302, DOI: 10.3892/ijo.2016.3467 (2016).

19. Mittler, F. et al. High-Content Monitoring of Drug Effects in a 3D Spheroid Model. Front. Oncol. 7, DOI: 10.3389/fonc.2017.00293 (2017).

20. Yang, W. et al. Mask-free generation of multicellular 3D heterospheroids array for high-throughput combinatorial anti-cancer drug screening. Mater. & Des. 183, 108182, DOI: 10.1016/j.matdes.2019.108182 (2019).

21. Seleci, D. A., Seleci, M., Stahl, F. & Scheper, T. Tumor homing and penetrating peptide-conjugated niosomes as multi-drug carriers for tumor-targeted drug delivery. RSC Adv. 7, 33378–33384, DOI: 10.1039/C7RA05071B (2017).

22. Zoetemelk, M., Rausch, M., Colin, D. J., Dormond, O. & Nowak-Sliwinska, P. Short-term 3D culture systems of various complexity for treatment optimization of colorectal carcinoma. Sci. Reports 9, 7103, DOI: 10.1038/s41598-019-42836-0 (2019).

23. Drexler, W. & Fujimoto, J. G. (eds.) Optical Coherence Tomography: Technology and Applications (Springer International Publishing, 2015), 2 edn.

24. Huang, Y. et al. Optical Coherence Tomography Detects Necrotic Regions and Volumetrically Quantifies Multicellular Tumor Spheroids. Cancer Res. 77, 6011–6020, DOI: 10.1158/0008-5472.CAN-17-0821 (2017).

25. Yan, F. et al. Characterization and quantification of necrotic tissues and morphology in multicellular ovarian cancer tumor spheroids using optical coherence tomography. Biomed. Opt. Express 12, 3352–3371, DOI: 10.1364/BOE.425512 (2021).

26. Gil, D. A., Deming, D. A. & Skala, M. C. Volumetric growth tracking of patient-derived cancer organoids using optical coherence tomography. Biomed. Opt. Express 12, 3789–3805, DOI: 10.1364/BOE.428197 (2021).

27. Roberge, C. L., Wang, L., Barroso, M. & Corr, D. T. Non-Destructive Evaluation of Regional Cell Density Within Tumor Aggregates Following Drug Treatment. J. Vis. Exp. JoVE DOI: 10.3791/64030 (2022).

28. Apelian, C., Harms, F., Thouvenin, O. & Boccara, A. C. Dynamic full field optical coherence tomography: subcellular metabolic contrast revealed in tissues by interferometric signals temporal analysis. Biomed. Opt. Express 7, 1511–1524, DOI: 10.1364/BOE.7.001511 (2016).

29. Scholler, J. et al. Dynamic full-field optical coherence tomography: 3D live-imaging of retinal organoids. Light. Sci. & Appl. 9, 140, DOI: 10.1038/s41377-020-00375-8 (2020).

30. El-Sadek, I. A. et al. Optical coherence tomography-based tissue dynamics imaging for longitudinal and drug response evaluation of tumor spheroids. Biomed. Opt. Express 11, 6231–6248, DOI: 10.1364/BOE.404336 (2020).

31. Leung, H. M. et al. Imaging intracellular motion with dynamic micro-optical coherence tomography. Biomed. Opt. Express 11, 2768–2778, DOI: 10.1364/BOE.390782 (2020).

32. Münter, M. et al. Dynamic contrast in scanning microscopic OCT. Opt. Lett. 45, 4766–4769, DOI: 10.1364/OL.396134 (2020).

33. Kohlfaerber, T. et al. Dynamic microscopic optical coherence tomography to visualize the morphological and functional micro-anatomy of the airways. Biomed. Opt. Express 13, 3211–3223, DOI: 10.1364/BOE.456104 (2022).

34. El-Sadek, I. A. et al. Three-dimensional dynamics optical coherence tomography for tumor spheroid evaluation. Biomed. Opt. Express 12, 6844–6863, DOI: 10.1364/BOE.440444 (2021).

35. Abd El-Sadek, I. et al. Label-free drug response evaluation of human derived tumor spheroids using three-dimensional dynamic optical coherence tomography. Sci. Rep. 13, 15377, DOI: 10.1038/s41598-023-41846-3 (2023).

36. Li, E., Makita, S., Hong, Y.-J., Kasaragod, D. & Yasuno, Y. Three-dimensional multi-contrast imaging of in vivo human skin by Jones matrix optical coherence tomography. Biomed.Opt.Express 8, 1290–1305, DOI: 10.1364/BOE.8.001290 (2017).

37. Costa, E. C., Gaspar, V. M., Coutinho, P. & Correia, I. J. Optimization of liquid overlay technique to formulate heterogenic 3d co-cultures models. Biotechnol. Bioeng. 111, 1672–1685, DOI: 10.1002/bit.25210 (2014).

38. Mukomoto, R. et al. Oxygen consumption rate of tumour spheroids during necrotic-like core formation. Analyst 145, 6342–6348, DOI: 10.1039/D0AN00979B (2020).

39. Yakavets, I. et al. Advanced co-culture 3d breast cancer model for investigation of fibrosis induced by external stimuli: optimization study. Sci. Rep. 10, 21273, DOI: 10.1038/s41598-020-78087-7 (2020).

40. Charlier, C. et al. Tamoxifen and its active metabolite inhibit growth of estrogen receptor-negative MDA-MB-435 cells. Biochem. Pharmacol. 49, 351–358, DOI: 10.1016/0006-2952(94)00492-5 (1995).

41. Levin, E. R. Integration of the Extranuclear and Nuclear Actions of Estrogen. Mol. Endocrinol. 19, 1951–1959, DOI: 10.1210/me.2004-0390 (2005).

42. Yao, J., Deng, K., Huang, J., Zeng, R. & Zuo, J. Progress in the Understanding of the Mechanism of Tamoxifen Resistance in Breast Cancer. Front. Pharmacol. 11 (2020).

43. Chen, G., Liu, W. & Yan, B. Breast Cancer MCF-7 Cell Spheroid Culture for Drug Discovery and Development. J. cancer therapy 13, 117, DOI: 10.4236/jct.2022.133009 (2022).

44. Gudimchuk, N. B. & McIntosh, J. R. Regulation of microtubule dynamics, mechanics and function through the growing tip. Nat. Rev. Mol. Cell Biol. 22, 777–795, DOI: 10.1038/s41580-021-00399-x (2021).

45. Hohmann, T. & Dehghani, F. The Cytoskeleton—A Complex Interacting Meshwork. Cells 8, 362, DOI: 10.3390/cells8040362 (2019).

46. McIntosh, J. R., Grishchuk, E. L. & West, R. R. Chromosome-microtubule interactions during mitosis. Annu. Rev. Cell Dev. Biol. 18, 193–219, DOI: 10.1146/annurev.cellbio.18.032002.132412 (2002).

47. Vale, R. D. The Molecular Motor Toolbox for Intracellular Transport. Cell 112, 467–480, DOI: 10.1016/S0092-8674(03)00111-9 (2003).

48. Papin, J. A., Hunter, T., Palsson, B. O. & Subramaniam, S. Reconstruction of cellular signalling networks and analysis of their properties. Nat. Rev. Mol. Cell Biol. 6, 99–111, DOI: 10.1038/nrm1570 (2005).

49. Schindelin, J. et al. Fiji: an open-source platform for biological-image analysis. Nat. Methods 9, 676–682, DOI: 10.1038/nmeth.2019 (2012).

