## Supplementary figures. for "Label-free visualization and quantification of the drug-type-dependent response of tumor spheroids by dynamic optical coherence tomography"

In this supplementary material, we provide nine figures. Figures S1, S4, and S7 present the cross-sectional OCT intensity, LIV, and OCDSl images extracted at the locations of the white arrows in the en face images presented in Figs. 1, 3, and 5 of the main manuscript. The cross-sectional images show tendencies similar to those of the en face images. Figures S2 and S3 present the results for two additional spheroids obtained under each treatment condition of the control and TAM. Figures S5 and S6 present the results for two additional spheroids obtained under each treatment condition of PTX, and Figs. S8 and S9 present the results for additional spheroids treated with DOX. The additional results are similar to those presented in the main manuscript. The results of control and TAM- treated, PTX- treated, and DOX-treated spheroids on TD-3 and TD-6 (Figs. S3, S6, and S9, respectively) were recently published as a conference proceeding [1].

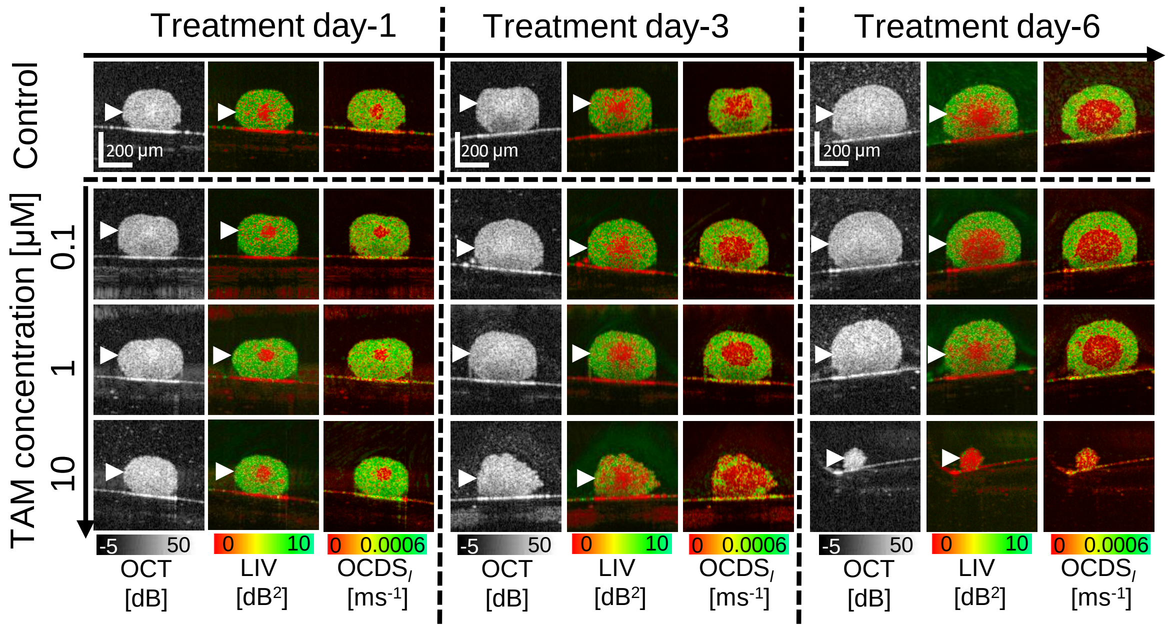


**Figure S1**: Cross-sectional OCT, LIV, and OCDS$_l$ images of the control and TAM-treated MCF-7 spheroids. The images are extracted at the white arrow head locations of Fig. 1 in the main manuscript.

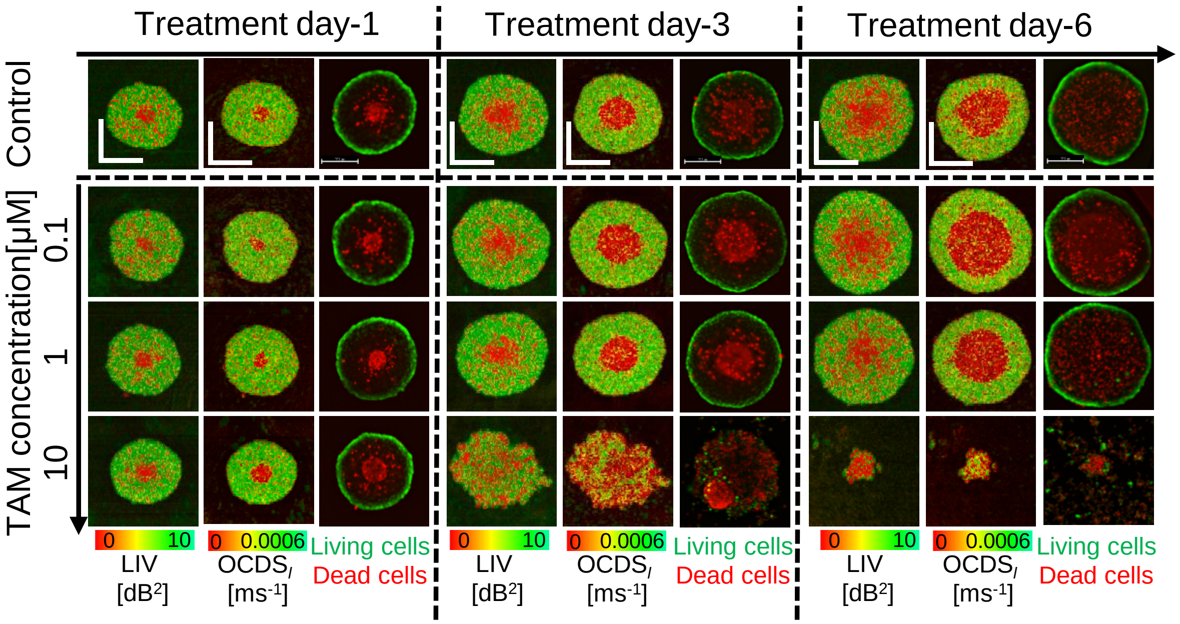


**Figure S2**: *En face* D-OCT and fluorescence images of additional control and TAM-treated MCF-7 spheroids measured under each treatment condition. The results are similar to those presented in Fig. 1 of the main manuscript. Scale bars represent 200 μm.

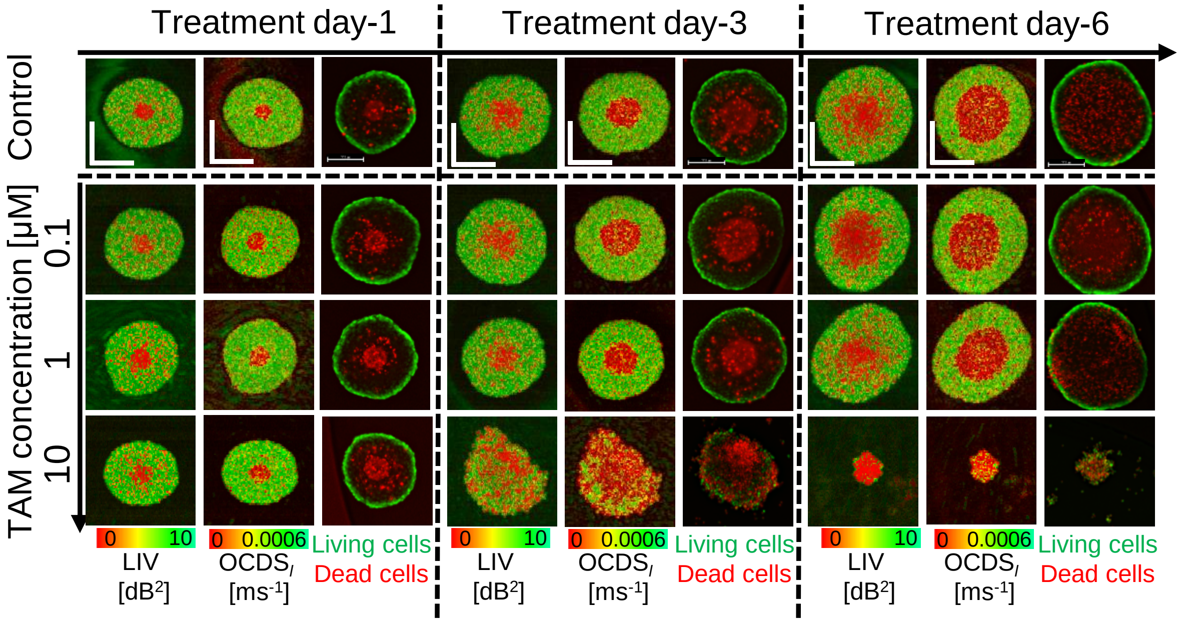


**Figure S3**: *En face* D-OCT and fluorescence images of another control and TAM-treated MCF-7 spheroids measured under each treatment condition. The results are similar to those presented in Fig. 1 of the main manuscript. Scale bars represent 200 μm.

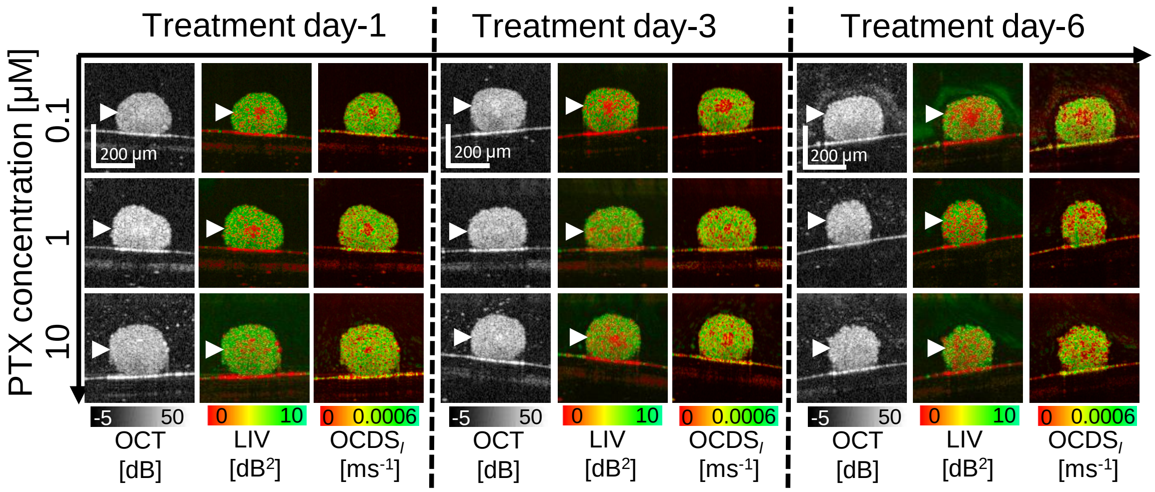


**Figure S4**: Cross-sectional OCT, LIV, and OCDS$_l$ images of the PTX-treated MCF-7 spheroids. The images are extracted at the white arrow head locations of Fig. 3 in the main manuscript.

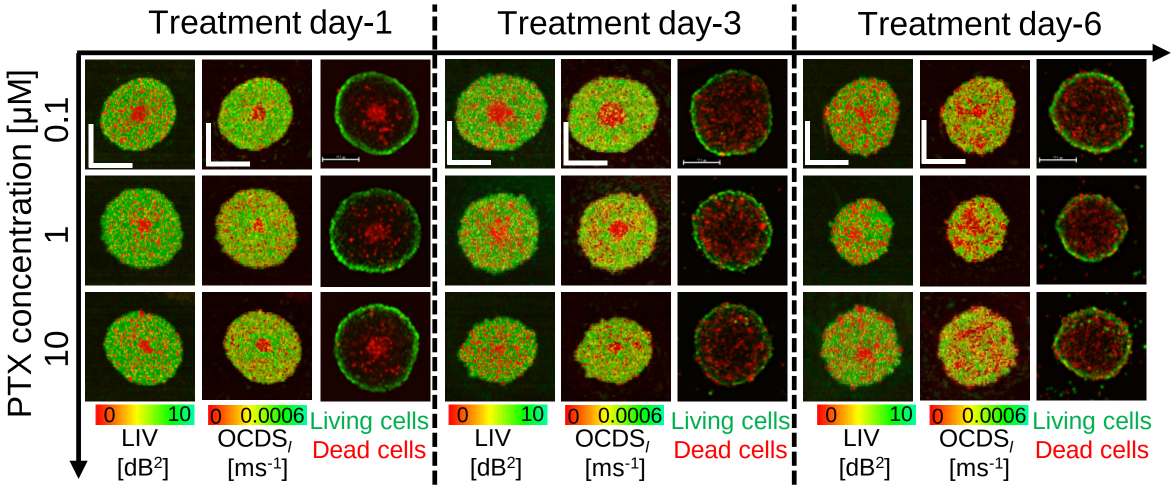


**Figure S5**: *En face* D-OCT and fluorescence images of additional PTX-treated MCF-7 spheroids measured under each treatment condition. The results are similar to those presented in Fig. 3 of the main manuscript. Scale bars represent 200 μm.

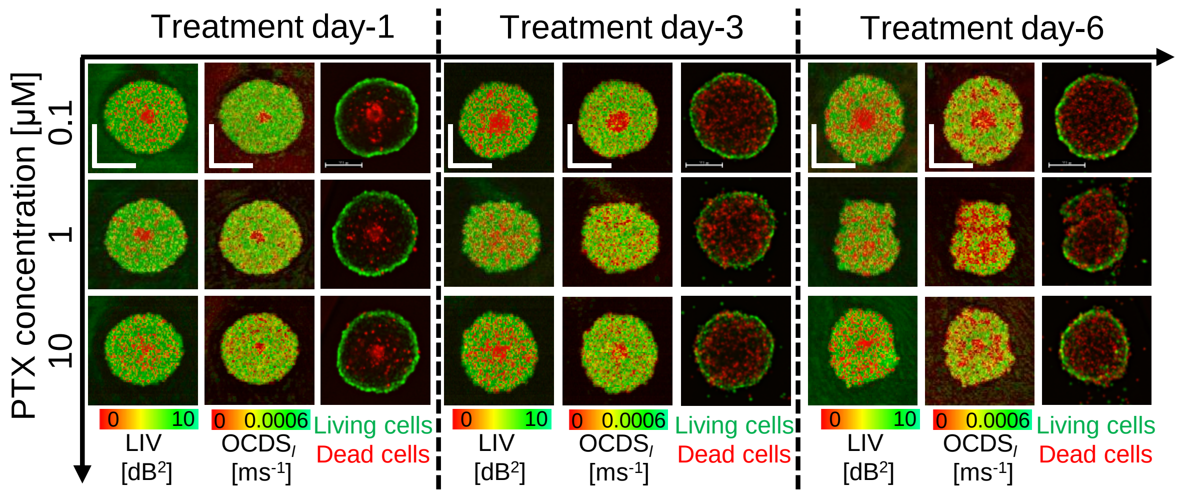


**Figure S6**: *En face* D-OCT and fluorescence images of another PTX-treated MCF-7 spheroids measure under each treatment condition. The results are similar to those presented in Fig. 3 of the main manuscript. Scale bars represent 200 μm.

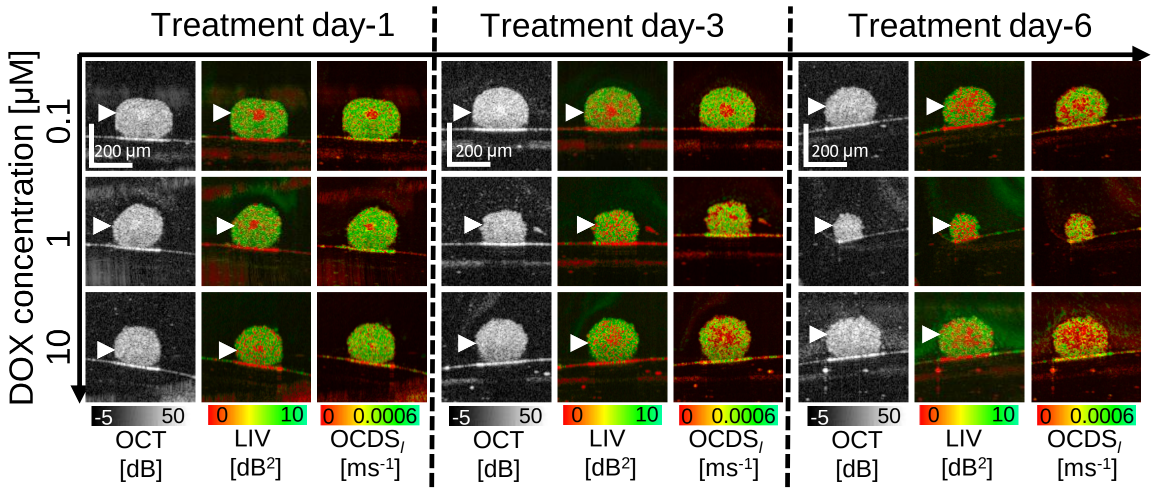


**Figure S7**: Cross-sectional OCT, LIV, and OCDS$_l$ images of the DOX-treated MCF-7 spheroids. The images are extracted at the white arrow head locations of Fig. 5 in the main manuscript.

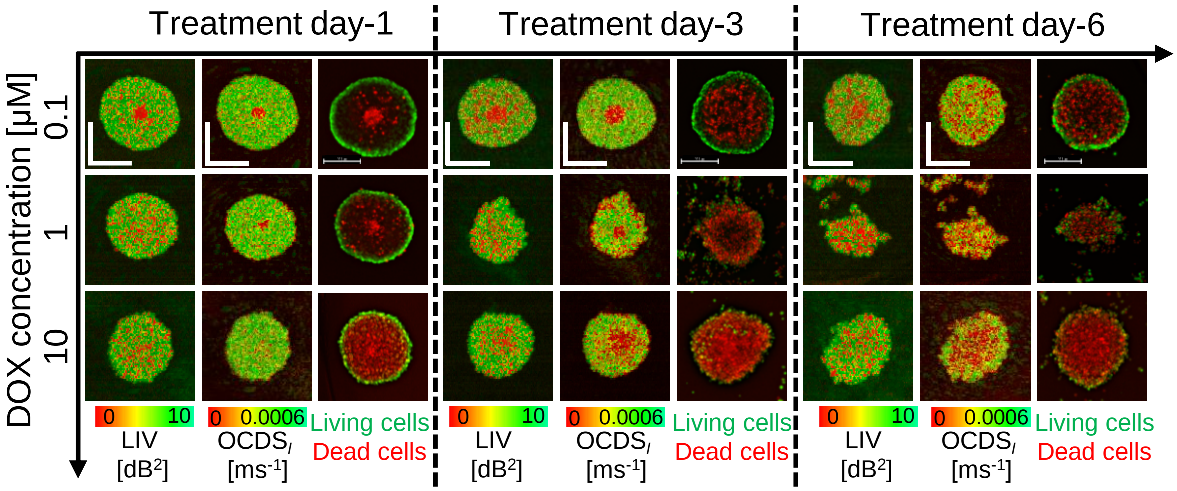


**Figure S8**: *En face* D-OCT and fluorescence images of additional DOX-treated MCF-7 spheroids measured under each treatment condition. The results are similar to those presented in Fig. 5 of the main manuscript. Scale bars represent 200 μm.

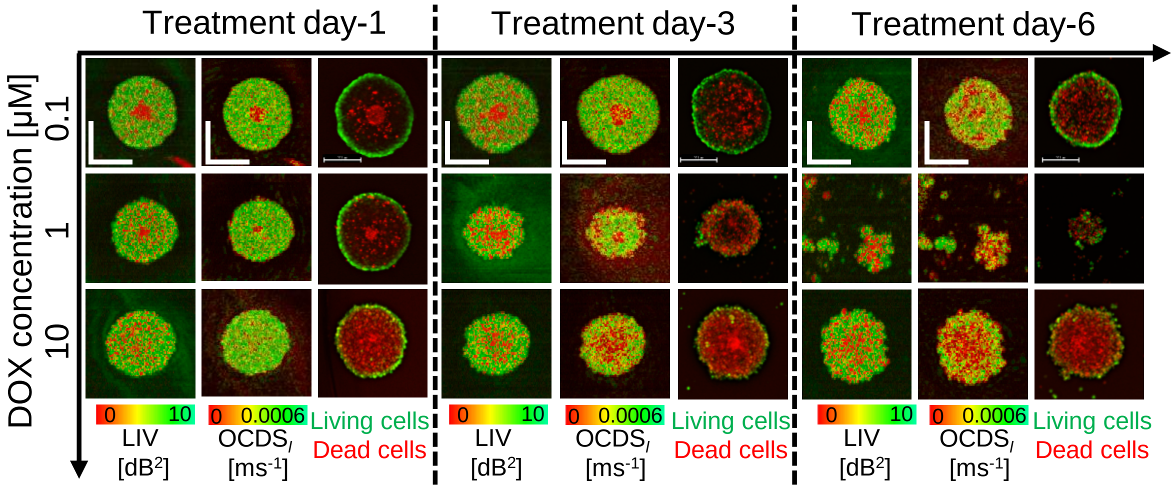


**Figure S9**: *En face* D-OCT and fluorescence images of another DOX-treated MCF-7 spheroids measured under each treatment condition. The results are similar to those presented in Fig. 5 of the main manuscript. Scale bars represent 200 μm.

### References


1. I. Abd El-Sadek, R. Morishita, T. Mori, S. Makita, P. Mukherjee, S. Matsusaka, and Y. Yasuno "Human-derived tumor-spheroid-based anti-cancer drugs testing using dynamic optical coherence tomography", *Proc. SPIE* 12632, Optical Coherence Imaging Techniques and Imaging in Scattering Media V, 126321A (11 August 2023).
